# Latent neural dynamics encode temporal context in speech

**DOI:** 10.1101/2021.07.15.452519

**Authors:** Emily P Stephen, Yuanning Li, Sean Metzger, Yulia Oganian, Edward F Chang

## Abstract

Direct neural recordings from human auditory cortex have demonstrated encoding for acoustic-phonetic features of consonants and vowels. Neural responses also encode distinct acoustic amplitude cues related to timing, such as those that occur at the onset of a sentence after a silent period or the onset of the vowel in each syllable. Here, we used a group reduced rank regression model to show that distributed cortical responses support a low-dimensional latent state representation of temporal context in speech. The timing cues each capture more unique variance than all other phonetic features and exhibit rotational or cyclical dynamics in latent space from activity that is widespread over the superior temporal gyrus. We propose that these spatially distributed timing signals could serve to provide temporal context for, and possibly bind across time, the concurrent processing of individual phonetic features, to compose higher-order phonological (e.g. word-level) representations.

## Introduction

Natural speech is a continuous stream of complex acoustic features, and listeners build representations of auditory objects at multiple levels, from phonemes, to syllables, words, and phrases (Berwick et al., 2013; Chomsky, 1985). The cortical basis of these dynamic compositional operations is an active area of research. There is evidence that the superior temporal gyrus (STG) performs speech-specific extraction of acoustic-phonetic features (Mesgarani et al., 2014), but where and how these segmental features are composed into longer units like words is less understood. Since the cascade of neural activity evoked by a given acoustic-phonetic feature can last longer than the feature itself (Gwilliams et al., 2020; Khalighinejad et al., 2017; Mesgarani et al., 2014; Näätänen and Picton, 1987; Norman-Haignere et al., 2020), there is potential for overlap in the neural representations over time. Hence the neural computations underlying speech comprehension should have a way to keep track of the temporal context of the individual phonetic units in order to compose them into a higher order unit such as a word (Fischer-Baum, 2018; Gwilliams et al., 2020).

We hypothesized that the mechanisms underlying temporal context tracking and composition in auditory cortex would be reflected in low-dimensional latent dynamics of electrocorticography (ECoG)-scale neural recordings. As neural recordings have grown in dimension, latent state models describing lower-dimensional summaries of populations of neurons have become more popular as the explanatory framework for understanding neural computation. In particular, there is a growing trend to map out geometric characteristics of latent states that could be indicative of the computational roles that are being played by the network (Russo et al., 2020, 2018; Seely et al., 2016; Vyas et al., 2020). One such geometrical motif is rotational dynamics (Churchland et al., 2012), which have been implicated in coordinating movements over time in the motor system (Buonomano and Laje, 2010; Cannon and Patel, 2021; Russo et al., 2020, 2018) (see Discussion). While the neural activity underlying speech perception is likely to be very different from that underlying motor sequencing, low-dimensional dynamics across the speech-responsive network in STG could reflect similar computational strategies to coordinate temporal context during speech perception.

There is already reason to believe that STG encodes information about timing: some STG populations respond to amplitude onset events found at the beginning of a sentence after silence period, or the acoustic edges that occur at the onset of vowels in syllables (called ‘peak rate’) (Hamilton et al., 2018; Oganian and Chang, 2019). If these signals are strong (representing a large proportion of the variance), temporally similar across different populations, and spatially widespread, they could constitute a low-dimensional latent state. In fact, Hamilton and colleagues (Hamilton et al., 2018) were able to find low-dimensional dynamics tied to sentence onsets using unsupervised linear dimensionality reduction. Unfortunately, due to the complex nature of the task (with a high-dimensional stimulus space and relevant stimulus features occurring closely in time), unsupervised methods have trouble uncovering dynamics related to other stimulus features, whose neural responses may overlap temporally and spatially with sentence onset responses. This makes it difficult to describe latent dynamics related to peak rate events, which are more closely aligned in timescale to the low-level compositional operations that we seek to describe. Supervised models, on the other hand, have historically focused on individual electrodes and as a result fail to describe latent dynamics that may reflect computational principles on a larger spatial scale.

Here we use a multivariate supervised approach to model the activity across all speech-responsive STG electrodes. Using integrative reduced rank regression (iRRR) (Li et al., 2019), we simultaneously estimate a separate low-dimensional latent state for each stimulus feature, including sentence onsets, peak rate events, and acoustic-phonetic features based on the place and manner of articulation. We find that iRRR outperforms models that treat each electrode individually, indicating that substantial feature-related information is shared across electrodes. The sentence onset and peak rate features explain more of the variance than phonetic features, reaffirming the importance of these timing-related features for encoding in STG. Furthermore, the latent states for the onset and peak rate are low-dimensional (5 and 6 dimensional, respectively) and distributed over centimeters of cortex, indicating a widespread signal that would be available to coordinate local and downstream processing. Geometrically, the latent dynamics contain a large proportion of rotational dynamics. Projections of the neural responses onto these low-dimensional spaces can be used to decode the time relative to the most recent sentence onset or peak rate event, with performance that is better than decoding from the full high-dimensional responses across all electrodes. We propose that the sentence onset response is an initialization signal and the peak rate latent states encode the time relative to acoustic events at the sentence and syllable scales. For peak rate, this spatially distributed timing signal could be used in local and downstream processing when composing word-level representations from low-level acoustic features.

## Results

### Model motivation and design

We modeled the high gamma (70-150 Hz) amplitude recorded on 331 speech-responsive electrodes located over the left superior temporal gyrus (STG) in 11 participants while they passively listened to 438 naturally spoken sentences from the Texas Instruments and Massachusetts Institute of Technology (TIMIT) acoustic-phonetic corpus (Garofolo et al., 1993). High gamma amplitudes in neural voltage recordings are known to correlate with the firing rates (Dubey and Ray, 2020; Manning et al., 2009; Ray et al., 2008; Ray and Maunsell, 2011; Scheffer-Teixeira et al., 2013) and dendritic processes (Bédard et al., 2006; Leszczyński et al., 2020; Miller et al., 2009; Suzuki and Larkum, 2017) of neurons near the electrode (Buzsáki et al., 2012), and we use them here as a proxy for the level of population activity under the ECoG electrodes. Using our model, we show that high gamma responses to speech stimuli across hundreds of electrodes can be parsimoniously represented as a combination of a few low-dimensional latent state responses to specific feature events in the stimulus. Two latent states in particular, corresponding to the sentence onset and peak rate features, reflect a large proportion of the explained variance in the model, and their dynamic properties suggest specific computational roles in the speech perception network.

Successful previous models of high gamma activity over STG have taken two different approaches: using supervised regression to model single-electrode responses as a function of spectral or linguistic characteristics in the audio speech signal (Aertsen and Johannesma, 1981; Holdgraf et al., 2017; Mesgarani et al., 2014; Oganian and Chang, 2019; Theunissen et al., 2001), and using unsupervised dimensionality reduction to infer latent states without reference to the characteristics of the audio stimulus (Hamilton et al., 2018).

The advantage of the single-electrode regression models is that they characterize the relationship between the neural responses and acoustic features in the speech signal. In the models, the high gamma responses on individual electrodes are considered to be the result of a convolution of time-dependent receptive fields with corresponding time series of acoustic features. The classic spectrotemporal receptive field (STRF) model, for example, uses a mel spectrogram of the stimulus as the acoustic feature representation, resulting in a framework where the neural receptive fields act as a linear filter on the speech spectrogram (Theunissen et al., 2001). Based on the observation that electrode activity over STG reflects information at the level of phonetic features rather than individual phonemes (Mesgarani et al., 2014), Oganian and Chang (Oganian and Chang, 2019) used an event-based feature representation to capture these effects and to show that some electrodes additionally have responses triggered by sentence onsets and sharp transients in the acoustic envelope of the speech signal, called peak rate events. While these models have been instrumental in describing the response patterns on individual electrodes, they fail to capture latent dynamics that are shared across multiple electrodes, which could uncover computational principles at work at a larger spatial scale.

An alternative approach uses unsupervised dimensionality reduction to investigate latent structure in neural responses to speech (e.g. (Hamilton et al., 2018)). Using convex nonnegative matrix factorization, they showed that electrodes can be naturally classified into two groups, “onset” electrodes that have a short increase in high gamma activity at the onset of a sentence, and “sustained” electrodes that show increased high gamma activity throughout the stimulus. This observation is also apparent using principal component analysis, in which the first component has characteristic sustained profile, and the second component has the onset profile (See Supplementary Figure S1). Note that the high gamma signals are not intrinsically low-dimensional: 2 dimensions capture only 24% of the variance in speech responsive electrodes (comparable to 16.9% of the variance in all electrodes captured in the first two clusters of (Hamilton et al., 2018)) and 189 dimensions are necessary to capture 80% of the variance. This could be related to the high-dimensional nature of the task: in an unsupervised framework in which the system responds to stimulus features, the response dimensionality needs to be at least as high-dimensional as the task itself (Gao et al., 2017; Stringer et al., 2019). Furthermore, both of these components are time-locked to sentence onset, and it is difficult to connect them or higher components to other speech features, possibly because the dynamics related to other features are not orthogonal to the sentence-onset subspace or to each other. In particular, the dependence of the neural responses on the peak rate events is not apparent from this analysis, and a model that could capture latent dynamics related to peak rate would be valuable for describing population encoding of shorter timescales.

We chose to use a model that combines the advantages of the regression and dimensionality reduction approaches, using a multivariate integrative reduced rank regression model (iRRR) (Li et al., 2019) to estimate the latent dynamics attributed to each speech feature separately. This group-reduced-rank model partitions the expected neural activity into a separate latent state for each feature, choosing the best latent dimensionality for each feature while penalizing the total dimensionality across all features. The model uses a multivariate adaptation of the event-based regression framework of Oganian and Chang (Oganian and Chang, 2019). In matrix form, the model has the following structure:

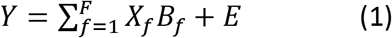

Where *Y* is the *T* × *N* matrix of high gamma amplitude values across electrodes and timepoints, each *X*_*f*_ (*T* × *D*) represents the delayed feature events for feature *f*, and *E* (*T* × *N*) is Gaussian noise, assumed to be uncorrelated across electrodes (*T*: number of timepoints; *N*: number of electrodes, *D*: number of delays, *F*: number of features). In *Y*, *X*, and *E*, the timepoints corresponding to subsequent sentence stimuli are stacked together. The coefficient matrices *B*_*f*_ (D × N) are the multivariate temporal response functions (MTRFs), representing the responses of each electrode to the given feature across electrodes and delays (up to 750ms).

Only speech responsive electrodes over STG were used for this analysis, defined using single-electrode fits to a linear spectrotemporal model (see electrode selection in Methods). Figure 1A shows the electrodes that were used, colored by the testing r^2^ value of the fitted spectrotemporal model: STG electrodes with r^2^>0.05 were used for subsequent analyses (*N* = 331).

**Figure 1:**
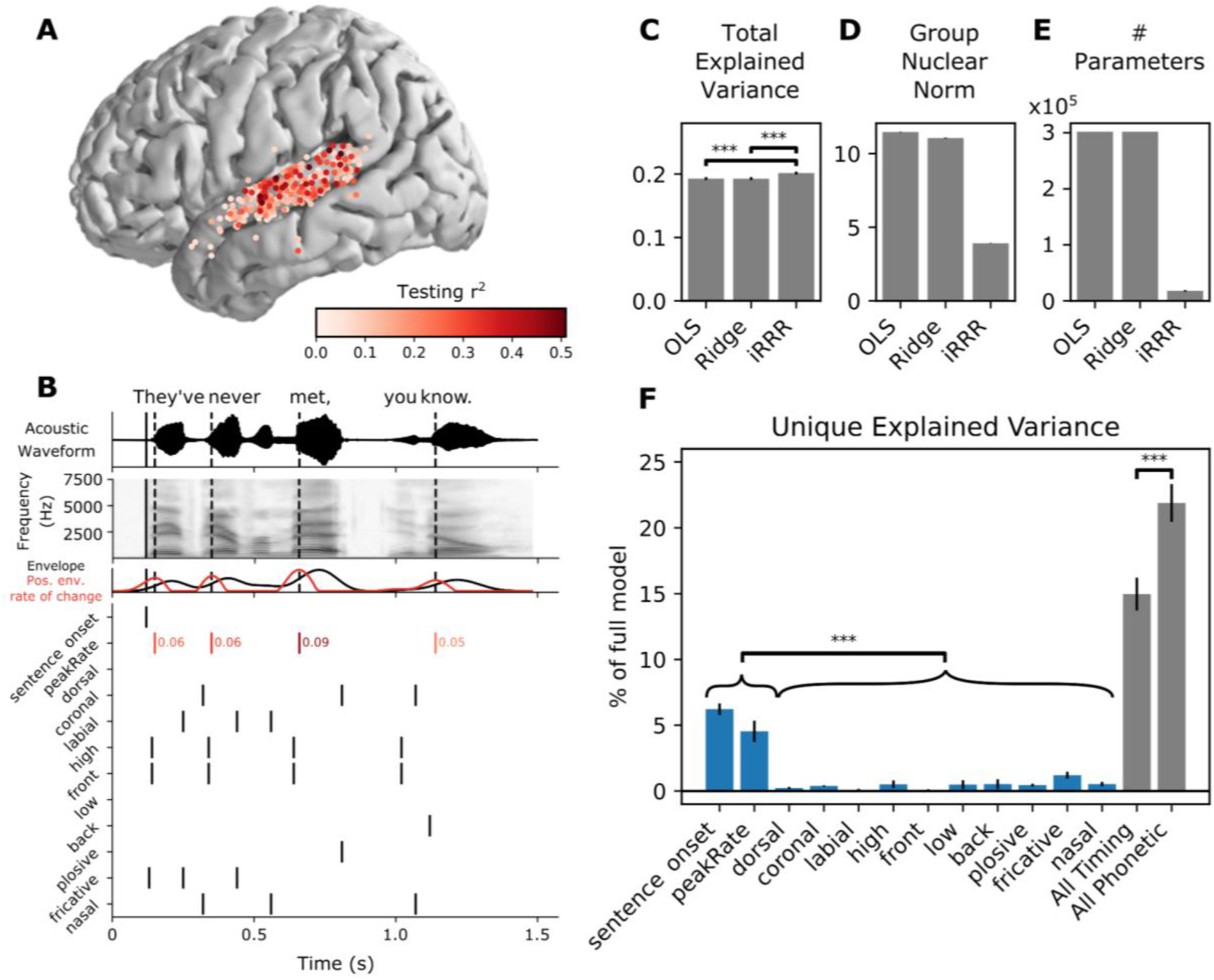
iRRR outperforms models that treat each electrode individually, and sentence onset and peak rate capture more of the variance than phonetic features. A: Electrodes used for model fitting, colored according to the testing r^2^ of the linear spectrotemporal (STRF) model (electrodes were selected for subsequent analysis if they were located over STG and if their testing r^2^ for the spectrotemporal model was greater than 0.05). B: Features used for feature temporal receptive field modeling. Top: the acoustic waveform of an example sentence. The solid vertical line shows the sentence onset event, and the dashed vertical lines show the times of the peak rate events. Second panel: the corresponding mel-band spectrogram. Third panel: the envelope of the acoustic waveform (black) and the positive rate of change of the envelope (red). The peaks in the positive envelope rate of change are the peak rate events. Bottom: the feature time series. White space represents no event (encoded by 0 in the feature matrix), black lines represent event times (encoded by 1), and red lines indicate peak rate event times with their corresponding magnitude indicated to the right. C, D, E: Performance of the iRRR model in comparison to ordinary least squares (OLS) and ridge regression (Ridge). 95% confidence intervals were estimated using the standard error of the mean across cross-validation folds (see Methods). Significance was assessed for comparisons using two-sided paired t-tests across cross-validation folds, *** p<0.0005. C: Total explained variance, computed as the testing r^2^ computed over all speech-responsive electrodes. D: Group nuclear norm, meaning the penalty term from the iRRR model (see Equation 2). E: The effective number of parameters for the fitted models. F: Unique explained variance for each feature (over all speech-responsive electrodes), expressed as a percentage of the variance captured by the full model. Comparing individual features, both timing features have significantly more unique explained variance than all phonetic features, after Bonferroni correction over pairs (left). Also shown is the unique explained variance for the combined timing features (sentence onset and peak rate) and the combined phonetic features (right). When the features are grouped, the phonetic features capture more unique explained variance than the timing features.

Figure 1B shows the feature events for an example sentence stimulus, “They’ve never met, you know”. The top two panels show the stimulus waveform and mel spectrogram, respectively, with the times of sentence onset and peak rate events indicated with vertical lines (solid and dashed, respectively). The features fall into two categories: timing (sentence onset and peak rate) and acoustic-phonetic (dorsal, coronal, labial, high, low, front, back, plosive, fricative, nasal). With the exception of peak rate, all of the feature events were encoded as binary time series with a 1 representing an event occuring, and 0 otherwise. For peak rate, the time series contained continuous values representing the slope of the acoustic amplitude signal at the time of maximal change, and 0 at all other times (in Figure 1B, red lines indicate peak rate event times and red numbers indicate the peak rate magnitude). We chose to include magnitude for peak rate events, because it is known to correlate very well with stressed syllables, i.e. syllables with higher stress will have higher peak rate magnitude.

We fit the regression model using integrative reduced-rank regression (iRRR) (Li et al., 2019), which applies a penalty based on a weighted sum of the nuclear norms of the feature matrices (see Methods for more detail):

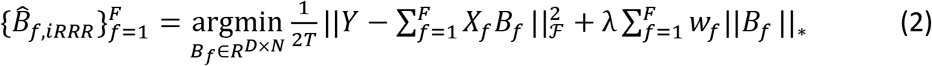

where || ⋅ ||_ℱ_ represents the Frobenius (L2) norm, the *w*_*f*_ s are chosen to balance the regularization across features, and λ is a regularization parameter. The notation || ⋅ ||^∗^ represents the nuclear norm, or the sum of the singular values of the bracketed matrix. The nuclear norm penalty acts as an L1 penalty on the singular values of each feature matrix, so the regression tends to find solutions where the feature matrices are low-rank (i.e. sparse in the singular values). Because many of the singular values will be zero, the fitted feature matrices can be represented using a low-dimensional singular value decomposition:

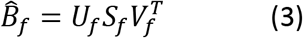

where *U*_*f*_ is *D* × *k*, *S*_*f*_ is *k* × *k*, and 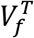 is *k* × *N*, for some *k* < *N*. In other words, the full multivariate feature receptive fields can be represented with a small number of patterns across time (columns of *U*_*f*_), patterns across electrodes (rows of 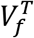), and corresponding weights (values on the diagonal of *S*_*f*_). The number of dimensions *k* can be different for each feature, and it comes from balancing the contribution of the feature to the first term (the mean squared error) with the contribution of the feature to the second term (the nuclear norm penalty), relative to other features. Increasing the tuning parameter λ will tend to increase the total number of dimensions used across all features.

For comparison, we also fit the same model using ordinary least squares (OLS) and ridge regression where a separate regularization parameter was chosen for each electrode. All models were fit using 10-fold cross validation. For the iRRR and ridge models, the regularization parameters were fit with an additional level of 5-fold cross-validation nested within the outer cross-validation.

### iRRR outperforms models that treat each electrode individually, and sentence onset and peak rate capture more of the variance than phonetic features

Figure 1C-E compare the three different fitting frameworks: OLS, ridge regression, and iRRR. Because the regression framework is the same for all three, the fitted models have very similar total explained variance (r^2^ computed over all electrodes, Figure 1C), but iRRR by design achieves a much smaller nuclear norm (Figure 1D), which results in solutions that can be described with 94% fewer parameters than OLS and ridge regression (Figure 1E). The fact that the iRRR model captures as much information as the single-electrode models using far fewer parameters suggests that substantial feature-related information is shared across electrodes.

In order to compare the contribution to the model of the different features, we fit reduced versions of the iRRR model with each feature left out. From there, we could compute the percent of explained variance by comparing the r^2^ of the full model 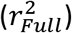 to the r^2^ of the model without feature 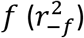:

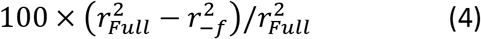

Figure 1F shows the result of this analysis: sentence onset and peak rate explain a larger percentage of the full model variance than each of the phonetic features (p<0.0005 for all comparisons using a two-sided paired t-test after Bonferroni correction). This suggests that these two timing features reflect a substantial amount of the speech-induced response across STG.

When the features are grouped into timing (sentence onset and peak rate) and phonetic (all other features) groups, both groups explain a large proportion of the variance (15% and 22%, respectively). Comparing the groups, however, the phonetic features explain more of the unique variance than the timing features (p<0.0005, two-sided paired t-test). This could be surprising in light of the individual feature comparisons: while timing features capture more explained variance than phonetic features when compared individually, when combined they capture less explained variance. This is likely due to (1) correlations between individual phonetic features that lead to lower individual unique explained variance and (2) the fact that more electrodes respond to sentence onset and peak rate than individual phonetic features (Oganian and Chang, 2019), meaning that sentence onset and peak rate have more widespread spatial support than the more spatially localized phonetic features. This more widespread spatial support means that the iRRR model is better able to consolidate the activity patterns across multiple electrodes, i.e. capture the latent dynamics, for the sentence onset and peak rate features than for the phonetic features. Accordingly, the following two sections describe the latent state representations for the sentence onset and peak rate features in more detail.

### The model fit captures known response differences between pSTG and mSTG

In Hamilton and colleagues’ (Hamilton et al., 2018) unsupervised model, the “onset” cluster of electrodes was found to occur primarily over the posterior portion of STG (pSTG). This observation led them to propose that pSTG may play a role in detecting temporal landmarks at the sentence and phrase level, because the short-latency, short-duration responses to sentence onsets in pSTG would be able to encode the event time with high temporal resolution. This idea fits well within a long history of evidence that stimulus responses in mSTG have longer latencies and longer durations than those in pSTG (Hamilton et al., 2020; Jasmin et al., 2019; Yi et al., 2019). Here, the model fits recapitulate these known differences between mSTG and pSTG.

As discussed above (Equation 3), the feature response matrices that are fitted by the iRRR model can be decomposed into a small number of components across time (“time components”, columns of *U*_*f*_), components across electrodes (“spatial components”, rows of 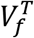), and corresponding weights (values on the diagonal of *S*_*f*_). Figure 2 shows the Sentence Onset and Peak Rate fitted feature matrices decomposed in this way (Since *U*_*f*_ and *V*_*f*_ are orthonormal, their columns are unit vectors: as a result, their units are arbitrary and can be best interpreted in relative terms).

**Figure 2:**
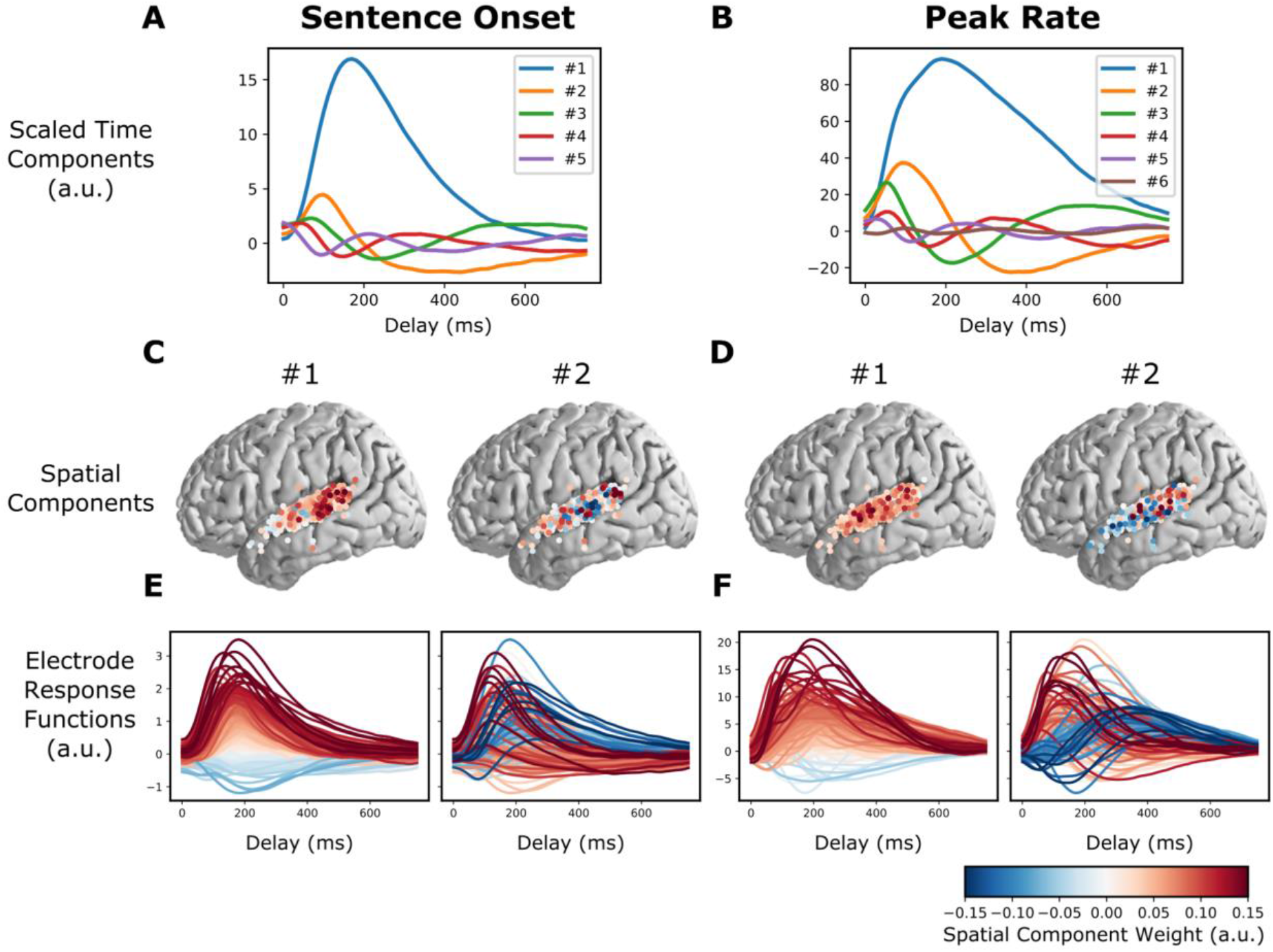
The model fit captures known response differences between pSTG and mSTG. A and B: Time components for the sentence onset and peak rate response matrices, scaled by their singular value (all panels of this figure use the fit from the first cross-validation fold). C: The first two spatial components (across electrodes) for sentence onset. E: The electrode responses to sentence onset events (rows of the sentence onset response matrix), colored by the first (left) or second (right) peak rate spatial component. The first spatial component for sentence onset shows that electrodes with large sentence onset responses (red lines in the left plot of E) tend to be in posterior STG (red circles in the left plot of C). D and F: (like C and E, but for peak rate). The second spatial component divides electrodes into fast and slow peak rate responses (red and blue lines in the right plot of F), which tend to occur over pSTG and mSTG, respectively (red and blue circles in the right plot of D).

Figures 2A and B show the time components scaled by their corresponding weights, and Figures 2C and D show the first two spatial components. To illustrate how the low dimensional components map back to the response functions for individual neurons, Figures 2E and F show the individual electrode response functions (rows of 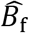), colored by the spatial component from Figures 2C and D.

Looking at the left panel of Figures 2C and 2E, we can see that electrodes that have large values in the first spatial component (red circles in Figure 2C, left) have relatively larger overall responses to sentence onset events (red lines in Figure 2E, left). These electrodes occur primarily over pSTG, which is in line with previous findings (Hamilton et al., 2018).

For peak rate, the first component plays the same role: electrodes that have larger values in the first spatial component (Figure 2D, left) have relatively larger overall responses to peak rate events (Figure 2F, left). Electrodes with large peak rate responses are not limited to pSTG like sentence onset electrodes: rather, they are distributed over all of STG. In other words, the encoding of peak rate in STG is not focal but is distributed over centimeters of cortex, suggesting a representation on a large spatial scale. Interestingly, the second component does appear to have a spatial distinction between pSTG and mSTG: electrodes with positive values for the second component tend to occur over pSTG, while electrodes with negative values for the second component tend to occur over mSTG (Figure 2D, right). The negative and positive values distinguish response functions by their temporal response profile: positive values correspond to electrodes that have an early peak rate response, while negative values correspond to electrodes that have a late peak rate response (Figure 2F, right). This suggests that peak rate responses over pSTG are faster than peak rate responses over mSTG.

### Feature latent states have rotational dynamics that capture continuous relative timing information

To show how the latent states behave during the presentation of a stimulus, we used the fitted model to predict the dynamics in each latent state during the presentation of the sentence “They’ve never met, you know” (Figure 3). Predictions from the model can be computed in latent space using the decomposition defined in Equation 3:

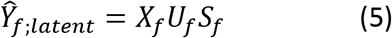

**Figure 3:**
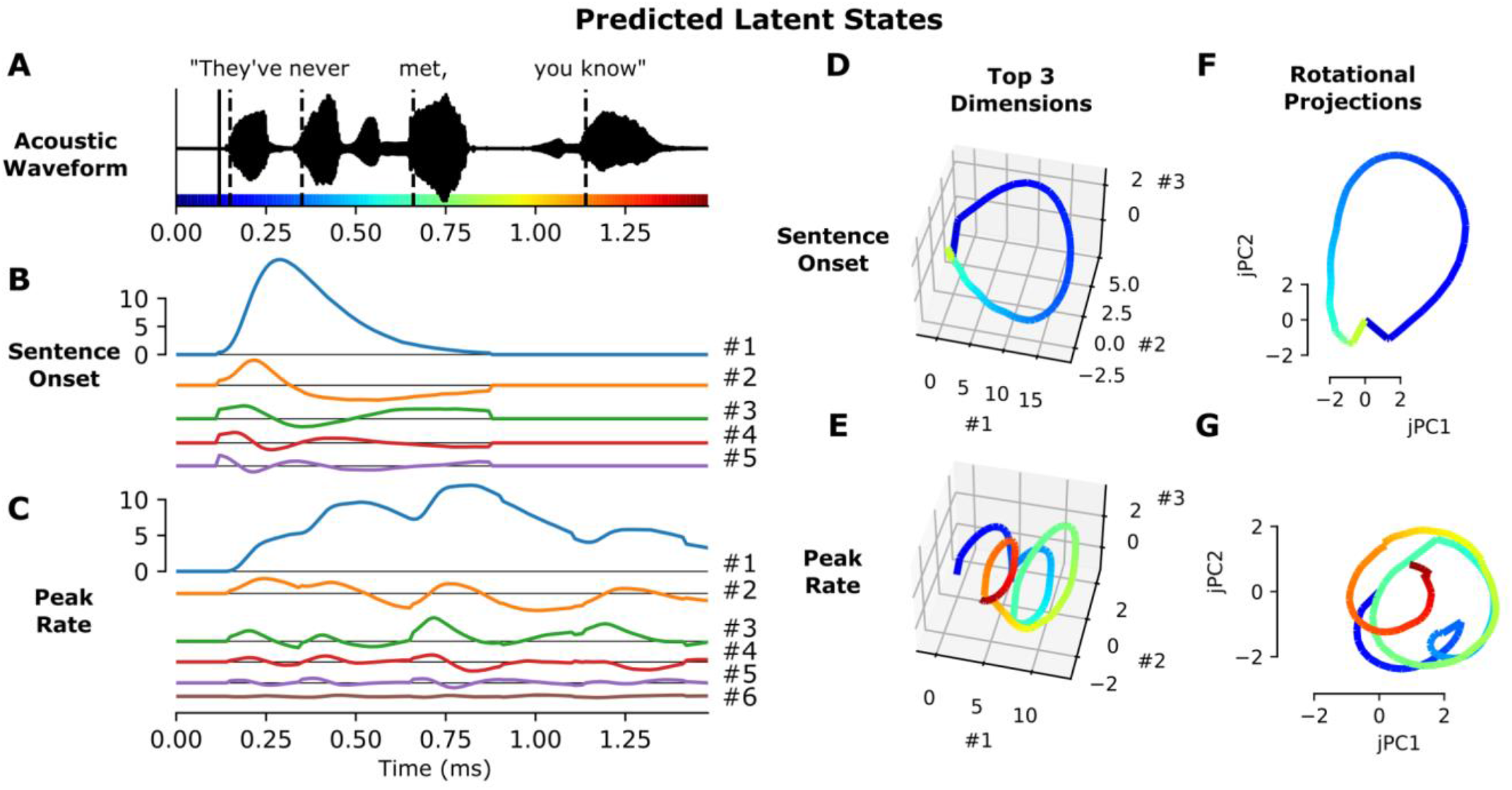
Feature latent states have rotational dynamics that capture continuous relative timing information. A: Acoustic waveform of the stimulus. Solid and dashed vertical lines indicate the timing of the sentence onset and peak rate events, respectively. Colors along the x-axis are used to indicate time parts D-G. B, C: Predicted latent states for the sentence onset and peak rate features corresponding to the given stimulus. D, E: Top three dimensions of the predicted sentence onset and peak rate latent states (the top three dimensions capture 98.7% and 98.8% of the variance in the sentence onset and peak rate coefficient matrices, respectively). F, G: Projection of the predicted sentence onset and peak rate latent states onto the plane of fastest rotation (identified using jPCA). The displayed jPCA projections capture 31.8% and 20.3% of the variance in the sentence onset and peak rate coefficient matrices, respectively. All panels of this figure use the fit from the first cross-validation fold.

The sentence onset latent space has 5 dimensions and the peak rate latent space has 6 dimensions. While the sentence onset feature only occurs once at the beginning of the stimulus, evoking a single response across the sentence onset dimensions, the peak rate feature occurs several times, and the dynamics of the peak rate latent state do not go back to baseline in between peak rate events (Figure 3B and C). Plotting the top three dimensions, which capture more than 98% of the variance in the coefficient matrices 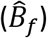, shows cyclical dynamics for both sentence onset and peak rate (Figure 3D and : the sentence onset state rotates once at the beginning of the sentence, and the peak rate latent state rotates 3-4 times, once after each peak rate event.

To quantify this effect, we used jPCA (Churchland et al., 2012) to identify the most rotational 2 dimensional subspace within the top three components of 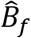. These planes capture 31.8% and 20.3% of the variance in the sentence onset and peak rate coefficient matrices, respectively, and they highlight the cyclical dynamics that were visible in the top 3 dimensions (Figure 3F and G).

Note that seeing cyclical dynamics in the latent states is not necessarily surprising: the coefficient matrices 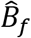 describe smooth multivariate evoked responses that will tend to start and end at the same baseline. We highlight them here to motivate a geometrical argument for the computational role of the peak rate responses (see Discussion) and to make the case that the structure of the peak rate responses enables them to act as a temporal context signal against which other features are organized. In order for the peak rate latent state to play this role, the trajectories should be sufficiently spread out in latent space to enable downstream areas to decode the time relative to the most recent peak rate event using just the instantaneous latent state. We investigate whether this is true in the next section.

### Latent states from the model can be used to decode time relative to feature events

So far, we have described how the model is fit using known feature event times, and how the fitted model can be used to predict responses given new feature events. We also wanted to know whether the model fit could be used to decode the timing of events, which would indicate that sufficient information is contained in the feature responses for downstream areas to use them as temporal context signals. The set of spatial components for each feature defines a feature-specific subspace of the overall electrode space. The projection of the observed high gamma time series onto this subspace is an approximation of the feature latent state (note that it is not exact, because the different feature subspaces are not orthogonal to each other):

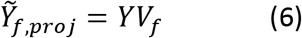

We asked whether this latent projection time series could be used to decode the time since the most recent feature event.

Figure 4 shows the result of this analysis: a perceptron model was trained to decode the time since the most recent feature event up to 750 ms, given either the activity on the full set of electrodes or the projection of the electrode activity onto the corresponding feature subspace (see Methods). The decoder for sentence onset performs slightly better when using all electrodes, which may be due to the large proportion of the overall activity that is time-locked to sentence onsets (see Supplementary Figure S1). For all other features, however, decoder performance using the reduced-dimensional latent subspaces performs even better than decoding using the full dimensional activity across electrodes (paired t-test over 10 cross validation folds, p<0.05 with Bonferroni correction across features). Because no information is gained in the projection operation, this is an indication that projecting onto the latent subspaces increases the signal to noise ratio, i.e. removes activity that is irrelevant to decoding relative time.

**Figure 4:**
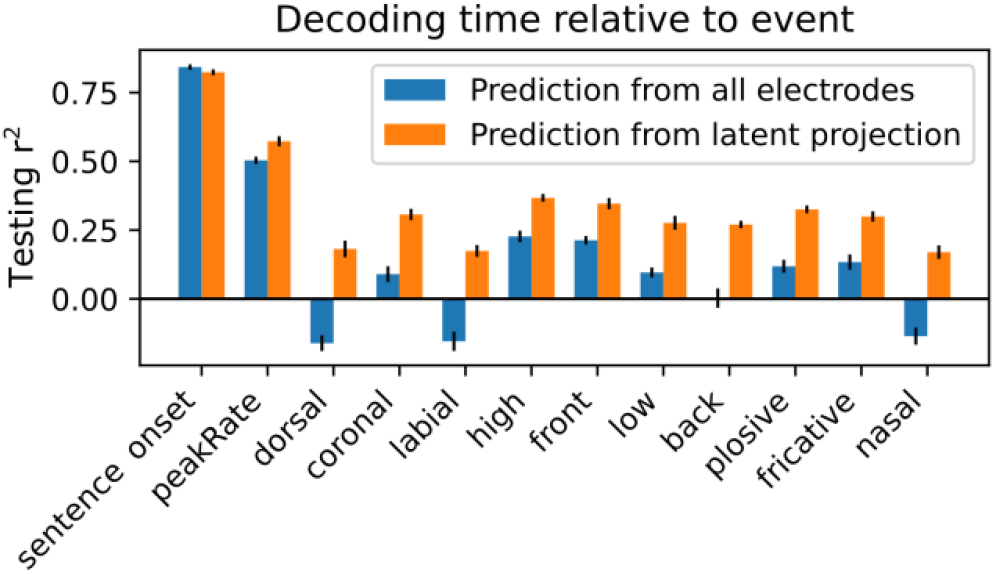
Latent states from the model can be used to decode time relative to feature events. Performance of a perceptron model trained to decode the time relative to the most recent feature event, for each feature. The models were trained either using the full high-dimensional set of high gamma responses across electrodes (blue bars) or using the projection of those responses onto the subspaces spanned by the feature latent states (orange bars). Performance is quantified using the testing set r^2^.

## Discussion

We have shown that a low dimensional regression model, iRRR, performs as well as classic models in representing high-gamma responses to timing and phonetic features of auditory stimuli, while using far fewer parameters. It accomplishes this compression by capturing similarities in feature responses that are shared across electrodes, which enables a low-dimensional latent state interpretation of the dynamics of high gamma responses to stimulus features. The sentence onset and peak rate features capture more unique variance than the other (phonetic) features, their responses are spread over both mSTG and pSTG, and their latent states show rotational dynamics that repeat after each event. Based on the geometry, duration, and spatial extent of the latent dynamics, we make the case that the sentence onset response could act as an initialization signal to kick the network into a speech-encoding state, while the peak rate response could provide a widespread temporal context signal that could be used to compose word-level representations from low-level acoustic and phonetic features.

The large magnitude of sentence onset responses in ECoG high gamma responses has been reported before (Hamilton et al., 2018): here, we confirm their large contribution to STG responses both using our iRRR model (Figure 1) and using PCA (Supplementary Figure S1). Importantly, the latent dynamics related to sentence onset last about 600 ms (Figure 2a). Since sentences in English often last longer than 600 ms (e.g. the sentences in the TIMIT corpus used here ranged from 900 ms to 2.6 s), these onset-related dynamics are unsuited to encode temporal context on an entire sentence level. Furthermore, sentence boundaries in continuous natural speech are rarely indicated with pauses or silence (Yoon et al., 2007), meaning that neural responses to acoustic onsets are unlikely to code sentence transitions. Rather, the latent dynamics in response to onsets may serve as a non-speech specific temporal indicator of the transition from silence to sound, occurring during perception of any auditory stimulus. During speech perception, the speech-related cortical networks could use this non-specific event as a reset or initialization signal. The idea that a large transient in the latent state could act to transition a network between states is also thought to occur in the motor system, where condition-invariant movement onset responses in the latent state mark the transition from motor preparation to motor behavior (Kaufman et al., 2016).

With regard to the peak rate dynamics, we propose that the computational role of the peak rate feature response is to keep track of word-level temporal context using a clock-like representation. The idea that structured latent state dynamics can act as clocks has been proposed in several different cognitive domains, most commonly in the motor system (Buonomano and Laje, 2010; Churchland et al., 2012; Remington et al., 2018; Vyas et al., 2020) (c.f. (Lebedev et al., 2020)) and in temporal interval estimation and perception (Cannon and Patel, 2021; Gámez et al., 2019; Mauk and Buonomano, 2004; Wang et al., 2018). In the motor system, Russo and colleagues (Russo et al., 2020) describe population dynamics in primary motor cortex (M1) and supplementary motor area (SMA) while a monkey performed a cyclic motor action. The population dynamics in M1 were rotational, exhibiting one rotation for each motor cycle, while the dynamics in SMA were shaped like a spiral, where 2-dimensional rotations for each motor cycle were translated along a third dimension. They proposed that this structure would be well-suited to keep track of progress through multi-cycle actions: each rotation encodes a single action, and translation along the third dimension encodes progress through the motor sequence. The rotational component of SMA population trajectories has also been suggested to operate as a time-keeping signal in auditory beat perception, where rotations through latent space keep track of the interval between beats (Cannon and Patel, 2021).

The peak rate latent state in STG could similarly be playing a computational role in auditory speech perception: the rotations in the peak rate subspace could serve to keep track of the time relative to the peak rate event, chunking time into intervals starting at the onset of a vowel. These intervals could then be used by downstream processing to give temporal context to the fine-grained phonetic feature information conveyed by other subpopulations. In other words, the rotational peak rate latent state could provide a temporal scaffolding on which individual phonetic features can be organized. Figure 5 illustrates this idea: when hearing the sentence “It had gone like clockwork,” the peak rate latent state partitions the sentence into four rotations, each one capturing the time since the most recent peak rate event. Downstream processing streams could combine this information with the phonetic feature information to put the phonetic feature events into their local context, here at the level of words or small sets of words (Figure 5C). Peak rate is in a unique position to play this role: it is the only feature that repeats within the linguistic structure of speech at the level of syllables/words, without reference to the linguistic contents. In addition, the peak rate responses are distributed over centimeters of cortex (Figure 2D) so the temporal context information would be widely available to local and downstream processing.

**Figure 5:**
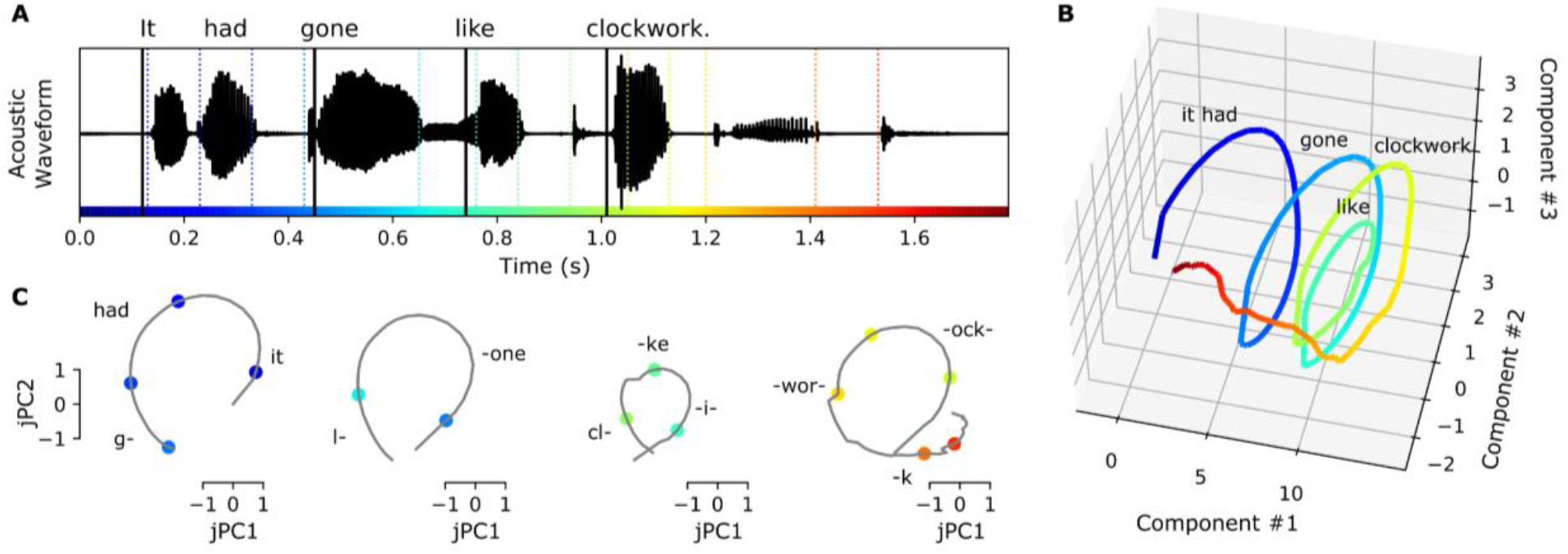
Peak rate rotational latent states could provide a temporal scaffolding on which individual acoustic features can be organized. A: The acoustic waveform for the stimulus “It had gone like clockwork”. Solid vertical lines indicate the times of peak rate events, and colored dashed vertical lines indicate the times of phonetic feature events. Colors are used to indicate time in all panels. B: The predicted peak rate latent state follows a spiral trajectory in the top 3 dimensions. C: Projected onto the plane of greatest rotation (jPC1 and 2), the predicted peak rate latent state divides the sentence into four intervals, each consisting of a rotation through state space that captures the time since the peak rate event occurred. Downstream processing could combine the relative time information encoded in the peak rate subspace (grey traces) with the feature identities encoded in the feature subspaces (colored points) to compose higher-order representations of words or small groups of words. Text in panels B and C indicates the approximate timing of the words in the stimulus.

In order for the peak rate latent state to play this role, it should have a couple of properties. First, there should be a mapping from points in state space to different relative times. As we showed in Figure 3, the rotational dynamics cause different relative times to be encoded in different locations of the latent space. Second, the trajectories in latent space should be consistent enough to support decoding of relative time in the presence of noise. In Figure 4, we showed that the projections of the neural activity onto the subspaces spanned by the feature latent states support decoding of the time relative to the most recent feature event. Note that while the latent state projections support decoding better than decoding from the full high-dimensional signal, the actual performance for peak rate is somewhat low (~50%). A possible reason for this could be that some peak rate events are more effective at driving the latent state than others (even after accounting for peak rate magnitude, as the model does), resulting in inconsistent decoding of the time since the most recent peak rate event.

Beyond the two-dimensional rotational dynamics, the peak rate latent trajectory forms a spiral in 3 dimensions (Figure 5B), similar to population trajectories in SMA during motor sequences (Russo et al., 2020). This suggests that the peak rate subpopulation may additionally encode the ordering of the word-level intervals within a larger linguistic context, such as the phrase level.

Furthermore, the representation of these intervals does not require top-down predictive coding (Hovsepyan et al., 2020; Lewis and Bastiaansen, 2015; Park et al., 2015; Pefkou et al., 2017) or entrainment of ongoing oscillations (Canolty, 2007; Ghitza, 2011; Giraud and Poeppel, 2012; Hovsepyan et al., 2020; Martin, 2020; Pittman-Polletta et al., 2020): in our model they are implemented via event-related potentials triggered by discrete acoustic (peak rate) events. While top-down and oscillatory mechanisms may play important roles in speech perception, our model demonstrates that some speech segmentation and context processing can be performed without them.

The events that we focus on for speech segmentation are peak rate events, moments of sharp increases in the acoustic envelope. The peak rate events in the model are coded with their magnitude (the slope of the rise in the acoustic envelope), which allows the model dynamics to change proportionally to the size of the event. This is important because peak rate events, also called auditory onset edges (Biermann and Heil, 2000; Doelling et al., 2014; Heil and Neubauer, 2001), differ in magnitude based on the stress level of the corresponding syllable (Oganian and Chang, 2019). This means that the dynamics triggered by peak rate events are sensitive to prosodic structure, both stressed syllables within words and stressed words within phrases. To investigate this further, it would be helpful to use a speech stimulus corpus with more complex prosodic structure than the TIMIT corpus used here.

In summary, our model (iRRR) represents STG high gamma responses to natural speech stimuli as a superposition of responses to individual phonetic and timing features, where each feature has a corresponding low-dimensional latent state that is shared across electrodes. It performs as well as single electrode models while using far fewer parameters, indicating that substantial feature-related information is shared across electrodes. Sentence onset and peak rate events, features representing timing at the sentence and syllable scales, capture more unique variance than phonetic features. The latent dynamics for sentence onset and peak rate contain information about the time since the most recent (sentence onset or peak rate) event, and the information is distributed across centimeters of cortex. We make the case that for peak rate, this relative timing information could play a role in composing word-level representations from low-level acoustic features, without requiring oscillatory or top-down mechanisms.

## Author Contributions

Conceptualization: E.P.S, E.F.C., Y.O., Y.L., S.M.; Data Curation: Y.O.; Formal Analysis: E.P.S., Y.O., Y.L.; Funding acquisition: E.F.C.; Supervision: E.F.C.; Resources: E.F.C.; Software: E.P.S., Y.O.; Visualization: E.P.S., Y.L., S.M., Y.O.; Writing: - original draft: E.P.S.; Writing - review & editing: E.P.S., E.F.C., Y.O., Y.L., S.M..

## Acknowledgements

This work was supported by grants from the NIH (R01-DC012379 and U01-NS117765 to EFC). This research was also supported by Bill and Susan Oberndorf, The Joan and Sandy Weill Foundation, and The William K. Bowes Foundation. The authors would also like to thank the members of the Chang lab at UCSF as well as James Hieronymus and Benjamin Antin for valuable feedback.

## Methods

### Participants

Participants included 11 patients (6M/5F, age 31 +/− 12 years) undergoing treatment for intractable epilepsy. As a part of their clinical evaluation for epilepsy surgery, high-density intracranial electrode grids (AdTech 256 channels, 4mm center-to-center spacing and 1.17mm diameter) were implanted subdurally over the left peri-Sylvian cortex. All procedures were approved by the University of California, San Francisco Institutional Review Board, and all patients provided informed written consent to participate. Data used in this study was previously reported in (Hamilton et al., 2018).

### Experimental Stimuli

Stimuli consisted of 499 English sentences from the TIMIT acoustic-phonetic corpus (Garofolo et al., 1993), spoken by male and female speakers with a variety of North American accents. Stimuli were presented through free-field Logitech speakers at comfortable ambient loudness (~70 dB), controlled by a custom MATLAB script. Participants passively listened to the sentences in 4 blocks, each lasting about 4 minutes. A subset of 438 sentences were selected for analysis that were heard once by all 11 subjects. The sentences had durations between 0.9 and 2.6s, with a 400ms intertrial interval.

### Neural recordings and electrode localization

Neural recordings were acquired at a sampling rate of 3051.8 Hz using a 256-channel PZ2 amplifier or 512-channel PZ5 amplifier connected to an RZ2 digital acquisition system (Tucker-Davis Technologies, Alachua, FL, USA).

Electrodes were localized by coregistering a preoperative T1 MRI scan of the individual subject’s brain with a postoperative CT scan of the electrodes in place. Freesurfer was used to create a 3d model of the individual subjects’ pial surfaces, run automatic parcellation to get individual anatomical labels, and warp the individual subject surfaces into the cvs_avg35_inMNI152 average template (Desikan et al., 2006; Fischl et al., 2004). More detailed procedures are described in (Hamilton et al., 2017).

### Preprocessing

For each electrode, the high gamma amplitude time series were extracted from the broadband neural recordings as follows (Hamilton et al., 2018; Oganian and Chang, 2019). First, the signals were downsampled to 400 Hz, rereferenced to the common average in blocks of 16 channels (blocks shared the same connector to the preamplifier), and notch filtered at 60, 120, and 180 Hz to remove line noise and its harmonics. These LFP signals were then filtered using a bank of 8 Gaussian filters with center frequencies logarithmically spaced between 70 and 150 Hz. Using the Hilbert transform, the amplitude of the analytic signal was computed for each of these frequency bands, and for each electrode the high gamma amplitude was defined as the first principal component across these 8 frequency bands. Finally, the high gamma amplitude was further downsampled to 100Hz and z-scored based on the mean and standard deviation across each experimental block.

### Electrode selection

In order select speech-responsive electrodes over STG, electrodes were included (1) if they were located over the STG, as identified in the Freesurfer anatomical parcellation of the individual subject cortical surface, and (2) if their high gamma activity was well-predicted by a linear spectrotemporal model (Hamilton et al., 2018).

For this single electrode analysis, the model had the form of a spectrotemporal receptive field (STRF):

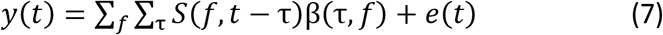

where *y* is the high gamma amplitude on a single electrode,*S* is the mel spectrogram of the speech audio signal over frequencies between 75Hz and 8kHz, coefficients β vary across frequencies and delays between 0 and 500ms, and *e* is the zero-mean Gaussian error term. Ridge regression was used to fit the models (see Model fitting below for details of the ridge regression framework): the data were split into 80% training and 20% testing data sets, the training data was used to choose the alpha parameter according to a 5-fold cross-validation, the full training data was fit using the chosen α parameter, and the r^2^ was assessed on the testing data (see Explained Variance Calculation below for computation of r^2^). Electrodes with r^2^>0.05 were included in subsequent analyses. The selected electrodes and their corresponding r^2^ values are shown in Figure 1A.

### Regression model setup

The multivariate temporal receptive field model has the following structure:

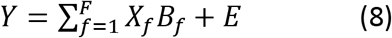

Where:

- *Y* is the *T* × *N* matrix of z-scored high gamma amplitude values across electrodes and timepoints. The time dimension represents a concatenation of all 438 sentence stimuli that were heard by every subject, from 500 ms before sentence onset until 500 ms after sentence offset (132,402 timepoints, later split for cross validation, see Model Fitting below). The electrode dimension includes speech-responsive electrodes from all subjects (331 electrodes).
- Each *X*_*f*_ (*T* × *D*) represents the delayed feature events for feature *f*. The first column contains the feature events across time (1 representing an event occuring, 0 otherwise. For peak rate, events were coded by a real-valued magnitude, see Figure 1B). Following columns contain the same time series, offset by time-delays between 10 ms and 750 ms (76 delays). There were 12 features: sentence onset, peak rate, dorsal, coronal, labial, high, front, low, back, plosive, fricative, and nasal (described below).
- *E* (*T* × *N*) is Gaussian noise, assumed to be uncorrelated across electrodes
- *B*_*f*_ (*D* × *N*) are the coefficient matrices, i.e. the multivariate temporal response functions (MTRFs), representing the responses of each electrode to the given feature across electrodes and delays
- *T*: number of timepoints; *N*: number of electrodes, *D*: number of delays, *F*: number of features.

Sentence onset was defined as the sound onset time for the sentence stimulus. Peak rate was extracted by taking the derivative of the analytic envelope of the speech signal: the peak rate event times were the times when the derivative reached a maximum, and the peak rate magnitude was the value of the derivative at that time point (Oganian and Chang, 2019). Phonetic feature event times (dorsal, coronal, labial, high, front, low, back, plosive, fricative, nasal) were extracted from time-aligned phonetic transcriptions of the TIMIT corpus, which were timed to the onset of the respective phonemes in the speech signal (Garofolo et al., 1993).

### Model fitting

The model was fit using ordinary least squares (OLS), ridge regression, and iRRR. The difference between the three is the objective function that is minimized to choose the fitted coefficient matrices:

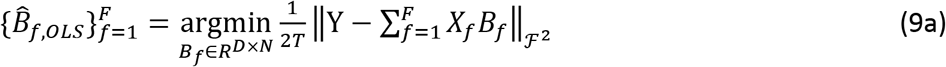

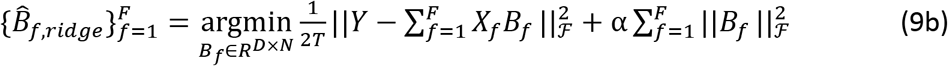

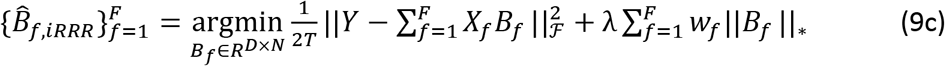

The weights used for the iRRR model were chosen to balance the different features (Li et al 2019):

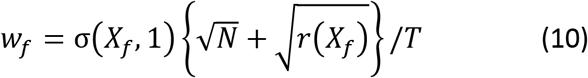

where σ(*X*_*f*_, 1) is the first singular value of the matrix *X*_*f*_ and *r*(*X*_*f*_) = *D* is the rank of matrix *X*_*f*_. All predictors *X*_*f*_ and responses *Y* were column-centered before fitting the models.

In order to compute confidence intervals for model performance metrics, models were fit using 10-fold cross validation, using group cross validation to keep time points corresponding to the same sentence stimulus in the same fold. For ridge regression and iRRR, an additional nested 5-fold cross validation was used to choose the α and λ parameters within each fold of the outer cross-validation.

Note that the approach of using a regression framework to fit a group-reduced rank model of neural activity has been used before (Aoi et al., 2020; Aoi and Pillow, 2019): the iRRR framework differs in that it uses an L1 relaxation, resulting in a convex optimization formulation that can be fit efficiently using alternating direction method of multipliers.

### Model performance metrics

Total explained variance (Figure 1C) was calculated as:

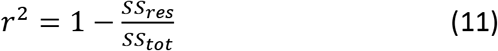

where the *SS*_*res*_ is the residual sum of squares computed on the testing dataset:

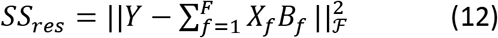

and *SS*_*tot*_ is the total sum of squares computed on the testing dataset:

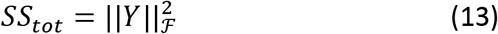

The group nuclear norm (Figure 1D) was computed as the penalty term in the iRRR model:

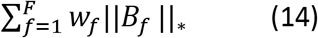

Because OLS and ridge regression yield full-rank coefficient matrices, the number of parameters (Figure 1E) used for both is *DN*. For iRRR, the number of parameters is *k*(*D* + *N* + 1), based on the singular value decomposition described in Equation 3, reproduced here:

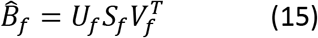

Unique explained variance for each feature (Figure 1F) was computed by fitting a reduced iRRR model without the feature *f*, and then comparing the total explained variance of the full model 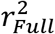 to the total explained variance of the reduced model 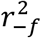. The reduced iRRR model was fit using the same λ value as the full model, chosen using nested cross validation on the full model as described above. For the “all timing” category, the reduced model was fit without sentence onset and peak rate, and for the “all phonetic” category, the reduced model was fit without the phonetic features. The unique explained variance was expressed as a percentage of the full model:

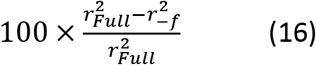

All metrics are reported in terms of the mean across the 10 folds of the cross validation, and 95% confidence intervals are 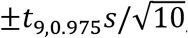, where *s* is the sample standard deviation across the 10 cross validation folds. Note that these confidence intervals do not account for the dependence between cross-validation folds due to reuse of samples in training and testing sets, and may therefore be smaller than the true intervals (Austern and Zhou, 2020; Bates et al., 2021; Bengio and Grandvalet, 2004).

Significant differences between conditions were assessed using paired two-tailed t-tests across cross-validation folds (Dietterich, 1998) for the following comparisons (with the resulting p-value ranges):

1. Total explained variance for OLS vs Ridge (p>0.05), OLS vs iRRR (p<0.0005), and Ridge vs iRRR (p<0.0005).
2. Unique explained variance of sentence onset vs each acoustic-phonetic feature and peak rate vs each acoustic-phonetic feature. Here the p-values were Bonferroni corrected across the (2 timing features times 10 acoustic-phonetic features) 20 comparisons. After correction, all comparisons were significant with p<0.0005.
3. Unique explained variance of the combined timing features vs the combined acoustic-phonetic features (p<0.0005).

Similar to the confidence intervals described above, the significance tests did not account for the dependence between cross-validation folds and may therefore have an inflated type II error (Austern and Zhou, 2020; Bates et al., 2021; Bengio and Grandvalet, 2004).

### Computing predicted responses

Given a fitted model, the predicted latent response to a stimulus matrix *X*_*f*_ is (reproduced from Equation 5):

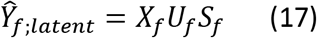

where, as before, *X*_*f*_ (*T* × *D*) represents the delayed feature events for feature *f*, *U*_*f*_ is the *D* × *k* time components for feature *f*, and *S*_*f*_ is a diagonal matrix containing the weights for each component 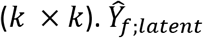 is a *T* × *k* matrix representing the predicted response within the *k*-dimensional latent space of the feature. Figure 3 shows the predicted sentence onset and peak rate responses to the sentence “They’ve never met, you know”.

### jPCA

The plane of fastest rotation for the sentence onset and peak rate latent states (Figure 3C) was identified by applying jPCA (Churchland et al., 2012) to the feature coefficient matrices 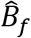. Using jPCA, we modeled the temporal receptive fields in the coefficient matrix as a linear dynamical system evolving over delays:

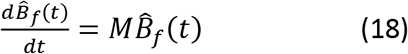

where *t* indexes the delay dimension of 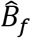, so the dynamical system describes the evolution of an *N*-dimensional dynamical system over *D* timepoints. By approximating the derivative on the left hand side using first differences, the transition matrix *M* can be fit using regression. Furthermore, the purely rotational component of the transition matrix can be isolated by constraining the matrix *M* to be skew-symmetric, having purely imaginary eigenvalues that come in complex conjugate pairs. The pair of eigenvectors with the largest magnitude eigenvalues describes the plane with the fastest rotations.

It is important to note that jPCA identifies planes with fast rotational dynamics, regardless of whether they capture a large proportion of the variance of the dynamics in the original dynamical system. Classic jPCA uses PCA in preprocessing in order to confine the analysis to six dimensions of largest variance. Here, the iRRR model chooses *k* dimensions for each feature that are most valuable to the overall fit of the model. Hence there was no need to perform additional PCA to reduce the dimensionality. However, because the coefficient matrices had dimensions capturing very little variance, we did subselect components to capture 98% of the variance of the coefficient matrices. For both sentence onset and peak rate, this corresponded to the top 3 components. Hence the jPCA plane represents the plane of maximal rotation within a 3-dimensional subspace capturing 98% of the variance in the 5-dimensional (or 6-dimensional) coefficient matrix for sentence onset (or peak rate). If we had used more components for the jPCA computation, the rotational dynamics would be stronger but they would capture much less of the variance (using *k* dimensions vs using 3 dimensions: 2.8% vs 31.8% for sentence onset and 4.8% vs 20.3% peak rate), making them less informative about the overall population dynamics.

Once the jPCs were computed using the coefficient matrices, the predicted trajectory for a given stimulus (Figure 3F and G) is calculated as:

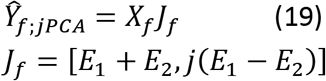

where *E*_1_ and *E*_2_ are the eigenvectors with largest eigenvalues of the skew-symmetric matrix *M* defined above. *J*_*f*_ is therefore the*N* × 2 projection matrix from electrode space onto the plane of highest rotation from jPCA.

### Event latency decoding

For the decoding analysis (Figure 4), a perceptron model was trained to predict the time relative to the most recent feature event (up to 750 ms). The model was designed using the MLPRegressor class of the sklearn package, with one hidden layer with 20 hidden units using a logistic activation function. We used a simple perceptron model in order to account for possible nonlinearities in the mapping from electrode space / feature latent space to relative times.

Using the same cross-validation framework that was used for iRRR model fitting, the perceptron model was trained using the training data (high gamma amplitudes) either across all electrodes *Y* or using the projected data onto the latent state subspace (reproduced from Equation 6):

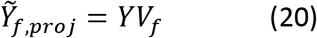

where *V*_*f*_ is the *N* × *k* matrix of electrode components for feature*f*, as above. The *T* × *k* matrix 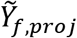 is an approximation of the latent state across time, but it may be contaminated by activity from other features because the *V*_*f*_ matrices do not describe orthogonal subspaces. It also contains activity from noise.

Performance of the models was assessed using r^2^ (Equation 11) on the held-out testing data for the cross-validation fold. The 95% confidence intervals were computed using the t distribution as described above, and the performance of the models trained on all-electrodes was compared to the performance of the models trained on the latent projections using a two-sided paired t test, as described above (Model Performance Metrics), Bonferroni corrected across the 12 features. The sentence onset model performed better using all electrodes than the latent projection (corrected p<0.05), while the models for all other features performed better using the latent projection than using all electrodes (corrected p<0.05).

### Code availability

Custom Python code to perform the iRRR fits is available online (https://github.com/emilyps14/iRRR_python), which is a port of the Matlab implementation by the original authors (https://github.com/reagan0323/iRRR, Li et al 2019). Python code for the analysis pipeline described above is also available (https://github.com/emilyps14/mtrf_python). We thank Antin and colleagues (Antin et al., 2021) for their implementation of jPCA in the Python programming language (https://github.com/bantin/jPCA), which we used to perform the jPCA.

## Supplementary Material

**Table S1.** Clinical and demographic details for subjects. Hem = hemisphere of implantation.

| Subject ID | Hem | Age | Sex | Handedness | Language dominance | Epilepsy focus |
| --- | --- | --- | --- | --- | --- | --- |
| SL01 | L | 44 | M | R | L | Left posterior STG |
| SL02 | L | 19 | M | R | L | Left anterior frontal lobe |
| SL03 | L |  | M |  | L | Left anterior temporal lobe |
| SL04 | L | 32 | F | R | L | Left anterior temporal lobe |
| SL05 | L | 25 | F | L | L | Left medial temporal lobe |
| SL06 | L | 31 | F | R | L | Left hippocampus/anterior lateral temporal |
| SL07 | L | 20 | F | R | L | Left hippocampus |
| SL08 | L | 60 | M | R (converted from L) | L | Left mesial temporal structures |
| SL09 | L | 26 | M | R | L | Left mesial and anterior lateral temporal cortex |
| SL10 | L | 22 | M | R | L | Left anterior temporal lobe |
| SL11 | L | 31 | F | R | L | Left hippocampus/amygdala |

**Figure S1:**
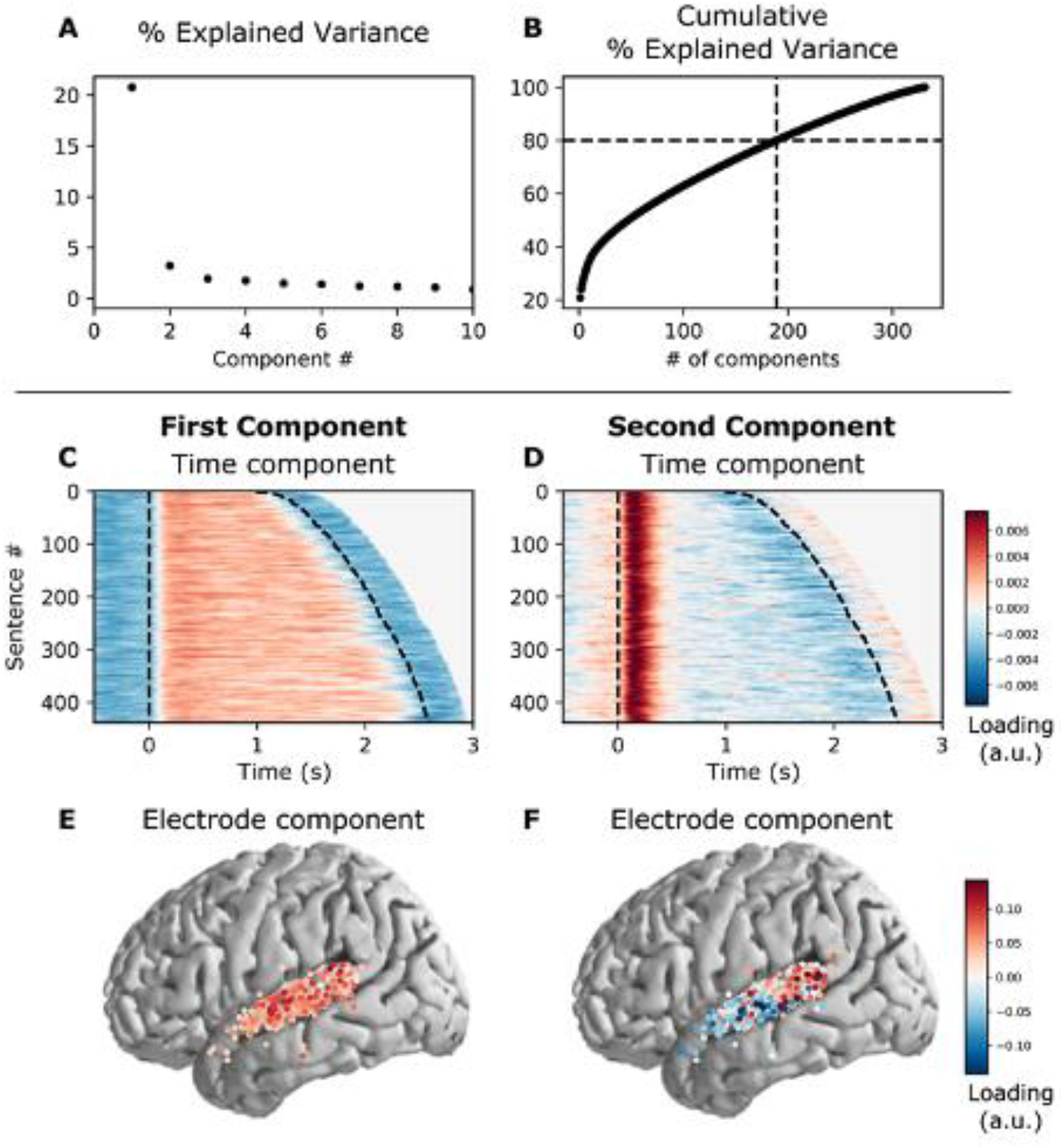
PCA partitions the high gamma activity across speech-responsive electrodes into a posterior onset response and a spatially widespread sustained response. A: The percent explained variance of the principal components. B: The cumulative percent explained variance. Note that 189 dimensions are required to capture 80% of the variance in the high gamma activity. C: The timecourse of the first component, aligned to sentence onset. Dashed lines indicate the start and end of the sentence stimulus, and the sentences have been ordered by their duration. This component has sustained responses, in the sense that the activity is high during the entire stimulus. D: The timecourse of the second component, aligned to sentence onset. This component has onset responses, in the sense that there is a short positive transient immediately after sentence onset. E: The spatial support of the first component. This component is spatially spread out over all of STG. F: The spatial support of the second component. This component is spatially divided, with strong positive weights over posterior STG.

## Bibliography

Aertsen, A.M.H.J., Johannesma, P.I.M., 1981. The Spectro-Temporal Receptive Field: A functional characteristic of auditory neurons. Biol. Cybern. 42, 133–143. https://doi.org/10.1007/BF00336731

Antin, B., Shenoy, K., Linderman, S., 2021. Probabilistic jPCA: a constrained model of neural dynamics., in: Cosyne Abstracts 2021. Presented at the Cosyne21, Online.

Aoi, M.C., Mante, V., Pillow, J.W., 2020. Prefrontal cortex exhibits multidimensional dynamic encoding during decision-making. Nat. Neurosci. 23, 1410–1420. https://doi.org/10.1038/s41593-020-0696-5

Aoi, M.C., Pillow, J.W., 2019. Model-based targeted dimensionality reduction for neuronal population data 15.

Austern, M., Zhou, W., 2020. Asymptotics of Cross-Validation. ArXiv200111111 Math Stat.

Bates, S., Hastie, T., Tibshirani, R., 2021. Cross-validation: what does it estimate and how well does it do it? ArXiv210400673 Math Stat.

Bédard, C., Kröger, H., Destexhe, A., 2006. Does the 1 / f Frequency Scaling of Brain Signals Reflect Self-Organized Critical States? Phys. Rev. Lett. 97, 118102. https://doi.org/10.1103/PhysRevLett.97.118102

Bengio, Y., Grandvalet, Y., 2004. No unbiased estimator of the variance of k-fold cross-validation. J. Mach. Learn. Res. 5, 1089–1105.

Berwick, R.C., Friederici, A.D., Chomsky, N., Bolhuis, J.J., 2013. Evolution, brain, and the nature of language. Trends Cogn. Sci. 17, 89–98. https://doi.org/10.1016/j.tics.2012.12.002

Biermann, S., Heil, P., 2000. Parallels between timing of onset responses of single neurons in cat and of evoked magnetic fields in human auditory cortex. J. Neurophysiol. 84, 2426–2439. https://doi.org/10.1152/jn.2000.84.5.2426

Buonomano, D.V., Laje, R., 2010. Population clocks: motor timing with neural dynamics. Trends Cogn. Sci. 14, 520–527. https://doi.org/10.1016/j.tics.2010.09.002

Buzsáki, G., Anastassiou, C.A., Koch, C., 2012. The origin of extracellular fields and currents — EEG, ECoG, LFP and spikes. Nat. Rev. Neurosci. 13, 407–420. https://doi.org/10.1038/nrn3241

Cannon, J.J., Patel, A.D., 2021. How Beat Perception Co-opts Motor Neurophysiology. Trends Cogn. Sci. 25, 137–150. https://doi.org/10.1016/j.tics.2020.11.002

Canolty, R.T., 2007. Spatiotemporal dynamics of word processing in the human brain. Front. Neurosci. 1, 185–196. https://doi.org/10.3389/neuro.01.1.1.014.2007

Chomsky, N., 1985. Syntactic structures, 14. printing. ed, Janua Linguarum Series minor. Mouton, The Hague.

Churchland, M.M., Cunningham, J.P., Kaufman, M.T., Foster, J.D., Nuyujukian, P., Ryu, S.I., Shenoy, K.V., 2012. Neural population dynamics during reaching. Nature 1–8. https://doi.org/10.1038/nature11129

Desikan, R.S., Ségonne, F., Fischl, B., Quinn, B.T., Dickerson, B.C., Blacker, D., Buckner, R.L., Dale, A.M., Maguire, R.P., Hyman, B.T., Albert, M.S., Killiany, R.J., 2006. An automated labeling system for subdividing the human cerebral cortex on MRI scans into gyral based regions of interest. NeuroImage 31, 968–980. https://doi.org/10.1016/j.neuroimage.2006.01.021

Dietterich, T.G., 1998. Approximate Statistical Tests for Comparing Supervised Classification Learning Algorithms. Neural Comput. 10, 1895–1923. https://doi.org/10.1162/089976698300017197

Doelling, K., Arnal, L., Ghitza, O., Poeppel, D., 2014. Acoustic landmarks drive delta-theta oscillations to enable speech comprehension by facilitating perceptual parsing. NeuroImage 85. https://doi.org/10.1016/j.neuroimage.2013.06.035

Dubey, A., Ray, S., 2020. Comparison of tuning properties of gamma and high-gamma power in local field potential (LFP) versus electrocorticogram (ECoG) in visual cortex. Sci. Rep. 10, 5422. https://doi.org/10.1038/s41598-020-61961-9

Fischer-Baum, S., 2018. A Common Representation of Serial Position in Language and Memory, in: Psychology of Learning and Motivation. Elsevier, pp. 31–54. https://doi.org/10.1016/bs.plm.2018.08.002

Fischl, B., van der Kouwe, A., Destrieux, C., Halgren, E., Ségonne, F., Salat, D.H., Busa, E., Seidman, L.J., Goldstein, J., Kennedy, D., Caviness, V., Makris, N., Rosen, B., Dale, A.M., 2004. Automatically Parcellating the Human Cerebral Cortex. Cereb. Cortex 14, 11–22. https://doi.org/10.1093/cercor/bhg087

Gámez, J., Mendoza, G., Prado, L., Betancourt, A., Merchant, H., 2019. The amplitude in periodic neural state trajectories underlies the tempo of rhythmic tapping. PLOS Biol. 17, e3000054. https://doi.org/10.1371/journal.pbio.3000054

Gao, P., Trautmann, E., Yu, B.M., Santhanam, G., Ryu, S., Shenoy, K., Ganguli, S., 2017. A theory of multineuronal dimensionality, dynamics and measurement 1–50. https://doi.org/10.1101/214262

Garofolo, J.S., Lamel, L.F., Fisher, W.M., Pallett, D.S., Dahlgren, N.L., Zue, V., Fiscus, J.G., 1993. TIMIT Acoustic-Phonetic Continuous Speech Corpus. https://doi.org/10.35111/17GK-BN40

Ghitza, O., 2011. Linking Speech Perception and Neurophysiology: Speech Decoding Guided by Cascaded Oscillators Locked to the Input Rhythm. Front. Psychol. 2. https://doi.org/10.3389/fpsyg.2011.00130

Giraud, A.-L., Poeppel, D., 2012. Cortical oscillations and speech processing: emerging computational principles and operations. Nat. Neurosci. 15, 511–517. https://doi.org/10.1038/nn.3063

Gwilliams, L., King, J.-R., Marantz, A., Poeppel, D., 2020. Neural dynamics of phoneme sequencing in real speech jointly encode order and invariant content (preprint). Neuroscience. https://doi.org/10.1101/2020.04.04.025684

Hamilton, L.S., Chang, D.L., Lee, M.B., Chang, E.F., 2017. Semi-automated Anatomical Labeling and Inter-subject Warping of High-Density Intracranial Recording Electrodes in Electrocorticography. Front. Neuroinformatics 11. https://doi.org/10.3389/fninf.2017.00062

Hamilton, L.S., Edwards, E., Chang, E.F., 2018. A Spatial Map of Onset and Sustained Responses to Speech in the Human Superior Temporal Gyrus. Curr. Biol. 28, 1860–1871.e4. https://doi.org/10.1016/j.cub.2018.04.033

Hamilton, L.S., Oganian, Y., Chang, E.F., 2020. Topography of speech-related acoustic and phonological feature encoding throughout the human core and parabelt auditory cortex. bioRxiv 2020.06.08.121624. https://doi.org/10.1101/2020.06.08.121624

Heil, P., Neubauer, H., 2001. Temporal Integration of Sound Pressure Determines Thresholds of Auditory-Nerve Fibers. J. Neurosci. 21, 7404–7415. https://doi.org/10.1523/JNEUROSCI.21-18-07404.2001

Holdgraf, C.R., Rieger, J.W., Micheli, C., Martin, S., Knight, R.T., Theunissen, F.E., 2017. Encoding and Decoding Models in Cognitive Electrophysiology. Front. Syst. Neurosci. 11. https://doi.org/10.3389/fnsys.2017.00061

Hovsepyan, S., Olasagasti, I., Giraud, A.-L., 2020. Combining predictive coding and neural oscillations enables online syllable recognition in natural speech. Nat. Commun. 11, 3117. https://doi.org/10.1038/s41467-020-16956-5

Jasmin, K., Lima, C.F., Scott, S.K., 2019. Understanding rostral–caudal auditory cortex contributions to auditory perception. Nat. Rev. Neurosci. 20, 425–434. https://doi.org/10.1038/s41583-019-0160-2

Kaufman, M.T., Seely, J.S., Sussillo, D., Ryu, S.I., Shenoy, K.V., Churchland, M.M., 2016. The Largest Response Component in the Motor Cortex Reflects Movement Timing but Not Movement Type. eneuro 3, ENEURO.0085-16.2016. https://doi.org/10.1523/ENEURO.0085-16.2016

Khalighinejad, B., Cruzatto da Silva, G., Mesgarani, N., 2017. Dynamic Encoding of Acoustic Features in Neural Responses to Continuous Speech. J. Neurosci. 37, 2176–2185. https://doi.org/10.1523/JNEUROSCI.2383-16.2017

Lebedev, M.A., Ninenko, I., Ossadtchi, A., 2020. Rotational dynamics versus sequence-like responses (preprint). Neuroscience. https://doi.org/10.1101/2020.09.16.300046

Leszczyński, M., Barczak, A., Kajikawa, Y., Ulbert, I., Falchier, A.Y., Tal, I., Haegens, S., Melloni, L., Knight, R.T., Schroeder, C.E., 2020. Dissociation of broadband high-frequency activity and neuronal firing in the neocortex. Sci. Adv. 6, eabb0977. https://doi.org/10.1126/sciadv.abb0977

Lewis, A.G., Bastiaansen, M., 2015. A predictive coding framework for rapid neural dynamics during sentence-level language comprehension. CORTEX 68, 155–168. https://doi.org/10.1016/j.cortex.2015.02.014

Li, G., Liu, X., Chen, K., 2019. Integrative multi-view regression: Bridging group-sparse and low-rank models. Biometrics 75, 593–602. https://doi.org/10.1111/biom.13006

Manning, J.R., Jacobs, J., Fried, I., Kahana, M.J., 2009. Broadband Shifts in Local Field Potential Power Spectra Are Correlated with Single-Neuron Spiking in Humans. J. Neurosci. 29, 13613–13620. https://doi.org/10.1523/JNEUROSCI.2041-09.2009

Martin, A.E., 2020. A Compositional Neural Architecture for Language. J. Cogn. Neurosci. 32, 1407–1427. https://doi.org/10.1162/jocn_a_01552

Mauk, M.D., Buonomano, D.V., 2004. THE NEURAL BASIS OF TEMPORAL PROCESSING. Annu. Rev. Neurosci. 27, 307–340. https://doi.org/10.1146/annurev.neuro.27.070203.144247

Mesgarani, N., Cheung, C., Johnson, K., Chang, E.F., 2014. Phonetic Feature Encoding in Human Superior Temporal Gyrus. Science 343, 1006–1010. https://doi.org/10.1126/science.1245994

Miller, K.J., Sorensen, L.B., Ojemann, J.G., Den Nijs, M., 2009. Power-law scaling in the brain surface electric potential. PLoS Comput. Biol. 5, e1000609. https://doi.org/10.1371/journal.pcbi.1000609.g005

Näätänen, R., Picton, T., 1987. The N1 Wave of the Human Electric and Magnetic Response to Sound: A Review and an Analysis of the Component Structure. Psychophysiology 24, 375–425. https://doi.org/10.1111/j.1469-8986.1987.tb00311.x

Norman-Haignere, S.V., Long, L.K., Devinsky, O., Doyle, W., Irobunda, I., Merricks, E.M., Feldstein, N.A., McKhann, G.M., Schevon, C.A., Flinker, A., Mesgarani, N., 2020. Multiscale integration organizes hierarchical computation in human auditory cortex (preprint). Neuroscience. https://doi.org/10.1101/2020.09.30.321687

Oganian, Y., Chang, E.F., 2019. A speech envelope landmark for syllable encoding in human superior temporal gyrus. Sci. Adv. 14.

Park, H., Ince, R.A.A., Schyns, P.G., Thut, G., Gross, J., 2015. Frontal Top-Down Signals Increase Coupling of Auditory Low-Frequency Oscillations to Continuous Speech in Human Listeners. Curr. Biol. 25, 1649–1653. https://doi.org/10.1016/j.cub.2015.04.049

Pefkou, M., Arnal, L.H., Fontolan, L., Giraud, A.-L., 2017. θ-Band and β-Band Neural Activity Reflects Independent Syllable Tracking and Comprehension of Time-Compressed Speech. J. Neurosci. 37, 7930–7938. https://doi.org/10.1523/JNEUROSCI.2882-16.2017

Pittman-Polletta, B.R., Wang, Y., Stanley, D.A., Schroeder, C.E., Whittington, M.A., Kopell, N.J., 2020. Differential contributions of synaptic and intrinsic inhibitory currents to speech segmentation via flexible phase-locking in neural oscillators (preprint). Neuroscience. https://doi.org/10.1101/2020.01.11.902858

Ray, S., Crone, N.E., Niebur, E., Franaszczuk, P.J., Hsiao, S.S., 2008. Neural Correlates of High-Gamma Oscillations (60-200 Hz) in Macaque Local Field Potentials and Their Potential Implications in Electrocorticography. J. Neurosci. Off. J. Soc. Neurosci. 28, 11526–11536. https://doi.org/10.1523/JNEUROSCI.2848-08.2008

Ray, S., Maunsell, J.H.R., 2011. Different Origins of Gamma Rhythm and High-Gamma Activity in Macaque Visual Cortex. PLoS Biol. 9, e1000610. https://doi.org/10.1371/journal.pbio.1000610.g008

Remington, E.D., Egger, S.W., Narain, D., Wang, J., Jazayeri, M., 2018. A Dynamical Systems Perspective on Flexible Motor Timing. Trends Cogn. Sci. 22, 938–952. https://doi.org/10.1016/j.tics.2018.07.010

Russo, A.A., Bittner, S.R., Perkins, S.M., Seely, J.S., London, B.M., Lara, A.H., Miri, A., Marshall, N.J., Kohn, A., Jessell, T.M., Abbott, L.F., Cunningham, J.P., Churchland, M.M., 2018. Motor Cortex Embeds Muscle-like Commands in an Untangled Population Response. Neuron 97, 953–966.e8. https://doi.org/10.1016/j.neuron.2018.01.004

Russo, A.A., Khajeh, R., Bittner, S.R., Perkins, S.M., Cunningham, J.P., Abbott, L.F., Churchland, M.M., 2020. Neural Trajectories in the Supplementary Motor Area and Motor Cortex Exhibit Distinct Geometries, Compatible with Different Classes of Computation. Neuron 107, 745–758.e6. https://doi.org/10.1016/j.neuron.2020.05.020

Scheffer-Teixeira, R., Belchior, H., Leão, R.N., Ribeiro, S., Tort, A.B.L., 2013. On high-frequency field oscillations (>100 Hz) and the spectral leakage of spiking activity. J. Neurosci. 33, 1535–1539. https://doi.org/10.1523/JNEUROSCI.4217-12.2013

Seely, J.S., Kaufman, M.T., Ryu, S.I., Shenoy, K.V., Cunningham, J.P., Churchland, M.M., 2016. Tensor Analysis Reveals Distinct Population Structure that Parallels the Different Computational Roles of Areas M1 and V1. PLOS Comput. Biol. 12, e1005164. https://doi.org/10.1371/journal.pcbi.1005164

Stringer, C., Pachitariu, M., Steinmetz, N., Carandini, M., Harris, K.D., 2019. High-dimensional geometry of population responses in visual cortex. Nature 571, 361–365. https://doi.org/10.1038/s41586-019-1346-5

Suzuki, M., Larkum, M.E., 2017. Dendritic calcium spikes are clearly detectable at the cortical surface. Nat. Commun. 8, 276. https://doi.org/10.1038/s41467-017-00282-4

Theunissen, F.E., David, S.V., Singh, N.C., Hsu, A., Vinje, W.E., Gallant, J.L., 2001. Estimating spatio-temporal receptive fields of auditory and visual neurons from their responses to natural stimuli. Netw. Comput. Neural Syst. 12, 289–316. https://doi.org/10.1080/net.12.3.289.316

Vyas, S., Golub, M.D., Sussillo, D., Shenoy, K.V., 2020. Computation Through Neural Population Dynamics. Annu. Rev. Neurosci. 43, 249–275. https://doi.org/10.1146/annurev-neuro-092619-094115

Wang, J., Narain, D., Hosseini, E.A., Jazayeri, M., 2018. Flexible timing by temporal scaling of cortical responses. Nat. Neurosci. 21, 102–110. https://doi.org/10.1038/s41593-017-0028-6

Yi, H.G., Leonard, M.K., Chang, E.F., 2019. The Encoding of Speech Sounds in the Superior Temporal Gyrus. Neuron 102, 1096–1110. https://doi.org/10.1016/j.neuron.2019.04.023

Yoon, T.-J., Cole, J., Hasegawa-Johnson, M., 2007. On the edge: Acoustic cues to layered prosodic domains, in: Proceedings of ICPhS. Citeseer, pp. 1264–1267.

